# PermaNet® Dual, a new deltamethrin-chlorfenapyr mixture net, shows improved efficacy against pyrethroid-resistant *Anopheles gambiae sensu lato* in southern Benin

**DOI:** 10.1101/2023.02.02.526745

**Authors:** Thomas Syme, Boris N’dombidjé, Martial Gbegbo, Damien Todjinou, Victoria Ariori, Patricia De Vos, Olivier Pigeon, Corine Ngufor

## Abstract

Pyrethroid-chlorfenapyr nets have demonstrated improved entomological and epidemiological impact in trials across Africa. This is driving increased demand for this novel net class in malaria endemic countries. PermaNet® Dual is a new deltamethrin-chlorfenapyr net developed by Vestergaard Sàrl to provide more options to malaria control programmes. We performed an experimental hut trial to evaluate the efficacy of PermaNet® Dual against wild, free-flying pyrethroid-resistant *Anopheles gambiae sensu lato* in Covè, Benin. PermaNet® Dual induced superior levels of mosquito mortality compared to a pyrethroid-only net and a pyrethroid-piperonyl butoxide net both when unwashed (77% with PermaNet® Dual vs. 23% with PermaNet® 2.0 and 56% with PermaNet® 3.0, p<0.001) and after 20 standardised washes (75% with PermaNet® Dual vs. 14% with PermaNet® 2.0 and 30% with PermaNet® 3.0, p<0.001). Using a provisional non-inferiority margin defined by the World Health Organisation, PermaNet® Dual was also non-inferior to a pyrethroid-chlorfenapyr net that has demonstrated improved public health value (Interceptor® G2), for vector mortality (79% vs. 76%, OR=0.854, 95% CIs: 0.703–1.038) but not for blood-feeding protection (35% vs. 26%, OR=1.445, 95% CIs: 1.203–1.735). PermaNet® Dual presents an additional option of this highly effective net class for improved control of malaria transmitted by pyrethroid-resistant mosquitoes.

## Background

Insecticide-treated nets (ITNs) are the most effective and widely adopted preventive measure against malaria. They have been consistently shown to reduce malaria morbidity and mortality under trial [1] and programmatic conditions [2], and have made the largest contribution of any intervention to recent reductions in malaria [3]. Their reliance however, on a single insecticide class – the pyrethroids – has exerted selective pressure favouring the spread of pyrethroid resistance in malaria vectors. Between 2010–2020, 88% of malaria-endemic countries detected pyrethroid resistance in at least one vector species [4]. Although studies show that ITNs remain protective against malaria infection despite resistance [5], a substantial body of evidence documents increased survival and blood-feeding of mosquitoes exposed to pyrethroid ITNs [6–9]. Given their importance in malaria prevention and control, any further loss in ITN effectiveness could contribute to resurgences in cases and deaths.

In response to this threat, dual-active ingredient ITNs combining a pyrethroid with another compound designed to restore control of pyrethroid-resistant malaria vectors have been developed. The first novel ITN type combines pyrethroids with piperonyl butoxide (PBO); a synergist that enhances pyrethroid efficacy by neutralising detoxifying enzymes associated with pyrethroid resistance [10]. Pyrethroid-PBO ITNs have shown improved entomological and epidemiological efficacy compared to pyrethroid-only ITNs in experimental hut [11–15] and cluster-randomised controlled trials (cRCTs) [16, 17]. They have since received a conditional recommendation from WHO for distribution in areas where vectors exhibit pyrethroid resistance leading to a significant increase in their deployment in endemic countries in recent years [18]. Pyrethroid-PBO ITNs are not however, without limitations. Notably, there are concerns over their durability following long-term household use [19]. Experimental hut trials in West Africa also suggest that pyrethroid-PBO ITNs may offer more limited benefits in areas with elevated pyrethroid resistance mediated by complex and multiple mechanisms [20]. More ITN types, ideally containing other novel insecticides to which vectors are susceptible are thus needed, for effective and sustainable vector control.

More recently, ITNs combining pyrethroids with chlorfenapyr, a pyrrole insecticide that disrupts mitochondrial function, have become available. Chlorfenapyr represents a new mode of action for public health which is suited for the control of vectors that have developed complex mechanisms of resistance to current insecticides. A pyrethroid-chlorfenapyr ITN developed by BASF (Interceptor® G2) has been prequalified by WHO [21], after demonstrating improved control of pyrethroid-resistant malaria vectors in experimental hut trials in Benin [22], Burkina Faso [23], Côte d’Ivoire [24] and Tanzania [25, 26]. Evidence of epidemiological impact is also emerging from large-scale trials and pilot distribution schemes in several countries. Most notably, cRCTs in Benin [27] and Tanzania [28] showed that Interceptor® G2 reduced child malaria incidence by 46% and 44% respectively over 2 years relative to standard pyrethroid-only ITNs. Pyrethroid-chlorfenapyr ITNs are soon expected to receive a WHO endorsement and policy recommendation, pending proof of improved public health impact from a second ongoing cRCT in Benin [29]. This is driving a substantial global increase in demand and order volumes for pyrethroid-chlorfenapyr ITNs for deployment in endemic countries [30]. The development of more innovative varieties of effective pyrethroid-chlorfenapyr nets from multiple manufacturers with robust production capacity, will help improve the health of the ITN market, increasing competition and leading to improved access to more affordable ITN products for optimal vector control impact [31].

PermaNet® Dual is a new deltamethrin-chlorfenapyr ITN developed by Vestergaard Sàrl. Recognising the prohibitive cost and time investment required to conduct cRCTs, to be prequalified by WHO and enter the ITN market successfully, PermaNet® Dual must be subjected to semi-field trials to establish its entomological superiority over standard pyrethroid-only ITNs. [32, 33]. It is also expected to demonstrate non-inferiority to a pyrethroid-chlorfenapyr ITN that has shown empirical evidence of improved public health value. To generate efficacy data as part of a PermaNet® Dual dossier submission for assessment by the Prequalification Unit Vector Control Product Assessment Team (PQT/VCP), we performed an experimental hut study to evaluate its efficacy and wash-resistance against wild, free-flying pyrethroid-resistant *Anopheles gambiae sensu lato* (*s.l*.) in Benin. PermaNet® Dual was tested unwashed and after 20 standardised washes and compared to three types of WHO prequalified ITNs; a pyrethroid-only net (PermaNet® 2.0), a pyrethroid-PBO net (PermaNet® 3.0) and a pyrethroid-chlorfenapyr net (Interceptor® G2). Data was analysed to assess the non-inferiority of PermaNet® Dual to Interceptor® G2 following a recent provisional WHO protocol [32]. The susceptibility of the vector population at the experimental hut site to the insecticides used in the ITNs was assessed during the trial using WHO bottle bioassays. Net pieces cut from ITNs before and after the hut trial were also tested in laboratory bioassays and analysed for chemical content. Following WHO PQT/VCP data requirements, the trial was performed in line with the Organisation for Economic Cooperation and Development (OECD) principles of good laboratory practice (GLP) at the CREC/LSHTM GLP-certified facility in Benin.

## Methods

### WHO bottle bioassays

WHO bottle bioassays [34] were performed using F1 progeny of field-collected *Anopheles gambiae s.l.* to assess the susceptibility of the vector population at the experimental hut station to the active ingredients used in the ITNs. Mosquitoes were exposed to the discriminating concentrations of deltamethrin (12.5 μg) and chlorfenapyr (100 μg). Additional exposures were performed with 2x (25 μg), 5x (62.5 μg) and 10x (125 μg) the discriminating concentration of deltamethrin to determine pyrethroid resistance intensity during the trial. To assess synergism and the contribution of cytochrome P450 monooxygenases to pyrethroid resistance, mosquitoes were also pre-exposed to PBO (400 μg) prior to deltamethrin-coated bottles (12.5 μg). Stock solutions for each insecticide were prepared by dissolving technical grade insecticide in acetone. Test bottles were coated by introducing 1 ml of stock solution into bottles and rotating manually. Approximately 100, unfed, 3–5-day old mosquitoes were exposed to each insecticide and dose for 60 mins in four batches of 25. Similar numbers of mosquitoes were concurrently exposed to acetone and PBO-coated bottles as controls. Knockdown was recorded after exposure and mosquitoes were transferred to labelled cups, provided access to 10% (w/v) glucose solution *ad libitum* and held at 27±2°C and 75±10% relative humidity (RH). Mortality was recorded after 24 h for deltamethrin and every 24 h up to 72 h for chlorfenapyr.

### Experimental hut trial

Experimental hut trials are standardised simulations of human-occupied housing designed to evaluate the efficacy of indoor vector control interventions against wild, free-flying mosquitoes under controlled field conditions. Host-seeking mosquitoes enter huts at night following attraction by odour cues emanating from human volunteers sleeping inside. Mosquitoes entering the huts then interact freely with the human host and vector control intervention and in the morning, they are collected and scored for physiological and behavioural parameters. The malaria control potential of vector control interventions is assessed primarily in terms of their ability to induce vector mortality (transmission control) and prevent blood-feeding (personal protection).

### Study site and experimental huts

The experimental hut trial was conducted at the CREC/LSHTM field station in Covè, southern Benin (7°14’N2°18’E). The site is located in a vast area of rice irrigation which provides extensive and permanent mosquito breeding sites. *An. coluzzii* and *An. gambiae sensu stricto* (*s.s*.) occur sympatrically with the former predominating. Recent studies show a high frequency and intensity of pyrethroid and organochlorine resistance but susceptibility to carbamates, organophosphates and pyrroles [35]. Genotyping and gene expression studies have revealed that pyrethroid resistance is mediated by a high frequency of the knockdown resistance (*kdr*) L1014F mutation and overexpression of cytochrome P450 monooxygenases [36]. Experimental huts used were of West African design, constructed from concrete bricks with cement-plastered walls, a corrugated iron roof and a polyethylene ceiling. Mosquitoes entered via four window slits with a 1 cm opening positioned on two sides of the hut. A wooden-framed veranda projected from the rear wall of each hut to capture exiting mosquitoes. Huts were surrounded by a water-filled moat to preclude mosquito predators.

### Experimental hut treatments

PermaNet® Dual, was compared to three other WHO pre-qualified ITNs; a pyrethroid-only net (PermaNet® 2.0), a pyrethroid-PBO net (PermaNet® 3.0), and a pyrethroid-chlorfenapyr net (Interceptor® G2). A description of the different ITNs tested in the trial is provided below.

- PermaNet® Dual (Vestergaard Sàrl) is a candidate 100-denier, polyester ITN coated with a combination of deltamethrin and chlorfenapyr at 2.1 g/kg and 5 g/kg respectively.
- Interceptor® G2 (BASF) is a WHO-prequalified 100-denier, polyester ITN coated with a combination of alpha-cypermethrin and chlorfenapyr at 2.4 g/kg and 4.8 g/kg respectively.
- PermaNet® 3.0 (Vestergaard Sàrl) is a WHO-prequalified ITN. The roof panel is made of 100-denier, polyethylene monofilament incorporating a combination of deltamethrin and PBO at 4 g/kg and 25 g/kg respectively. The side panels are made of 100-denier, polyester multifilament coated with deltamethrin at 2.1 g/kg.
- PermaNet® 2.0 (Vestergaard Sàrl) is a WHO-prequalified polyester ITN coated with deltamethrin at 1.4 g/kg.

An untreated polyester net developed to a similar technical specification as PermaNet® Dual was also tested as a negative control.

All ITNs were tested unwashed and washed 20 times as a proxy for insecticidal loss over 3 years of field use, as per WHO guidelines [37]. Nets were erected in huts by tying the four edges of the roof panel to nails positioned at the upper corners of hut walls. Nets were given 6 holes each measuring 4 x 4 cm to mimic wear-and-tear from routine use. Nine treatments arms were evaluated in nine experimental huts as follows:

1. Untreated polyester net (negative control)
2. PermaNet® 2.0 – unwashed (deltamethrin only)
3. PermaNet® 2.0 – washed 20x
4. PermaNet® 3.0 – unwashed (roof: deltamethrin plus PBO; sides: deltamethrin only)
5. PermaNet® 3.0 – washed 20x
6. Interceptor® G2 – unwashed (alpha-cypermethrin plus chlorfenapyr)
7. Interceptor® G2 – washed 20x
8. PermaNet® Dual – unwashed (deltamethrin plus chlorfenapyr)
9. PermaNet® Dual – washed 20x

### Experimental hut trial procedure

Human volunteers slept in huts between 21:00–06:00 to attract wild, free-flying mosquitoes. Each morning, volunteers collected all mosquitoes from the different compartments of the hut (under the net, room, veranda) using a torch and aspirator and placed them in labelled plastic cups. Mosquito collections were then transferred to the field laboratory for morphological identification and scoring of immediate mortality and blood-feeding. Surviving, female *An. gambiae s.l.* were provided access to 10% glucose (w/v) solution and held at ambient conditions. Delayed mortality was recorded every 24 h up to 72 h to account for the delayed action of chlorfenapyr. Mosquito collections were performed 6 days per week and on the 7th day, huts were cleaned and aired to prevent contamination before the next rotation cycle. Sleepers were rotated between huts daily while treatments were rotated weekly to mitigate the impact of variable host and hut positional attractiveness on mosquito entry. Six replicate nets were also used per treatment and rotated within the treatment daily. The trial continued for one full treatment rotation (9 weeks) between November 2020 and January 2021.

### Experimental hut trial outcome measures

The efficacy of the experimental hut treatments was expressed in terms of the following outcome measures:

1. **Hut entry** – number of female mosquitoes collected in experimental huts
2. **Deterrence (%)** – reduction in the number of mosquitoes collected in the treated hut relative to the untreated control hut. Calculated as follows:

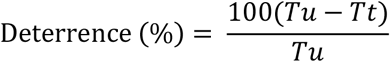

Where *Tu* is the number of mosquitoes collected in the untreated control hut and *Tt* is the number of mosquitoes collected in the treated hut.
3. **Exophily (%)** – exiting rates due to potential irritant effects of a treatment expressed as the proportion of mosquitoes collected in the veranda
4. **Blood-feeding (%)** – proportion of blood-fed mosquitoes
5. **Blood-feeding inhibition (%)** – proportional reduction in blood-feeding in the treated hut relative to the untreated control hut. Calculated as follows:

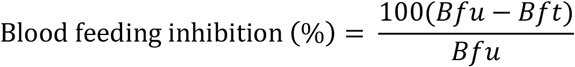

Where *Bfu* is the proportion of blood-fed mosquitoes in the untreated control hut and *Bft* is the proportion of blood-fed mosquitoes in the treated hut.
6. **Personal protection (%)** – reduction in the number of blood-fed mosquitoes in the treated hut relative to the untreated control hut. Calculated as follows:

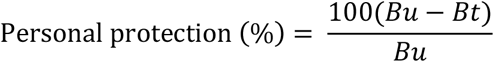

Where *Bu* is the number of blood-fed mosquitoes in the untreated control hut and *Bt* is the number of blood-fed mosquitoes in the treated hut.
7. **Delayed mortality (%)** – proportion of dead mosquitoes observed every 24 h up to 72 h after collection
8. **Overall killing effect (%)** – number of mosquitoes killed in the treated hut relative to the number collected in the untreated control hut. Calculated as follows:

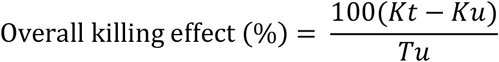

Where *Kt* is the number of dead mosquitoes in the treated hut, *Ku* is the number of dead mosquitoes in the untreated control hut and *Tu* is the number of mosquitoes collected in the untreated control hut.

### Preparation of net pieces for bioassays and chemical analysis

For each ITN type, a total of 5 net pieces (one from each panel) measuring 30 x 30 cm were cut before and after the hut trial from randomly selected unwashed and washed nets. Because of the mosaic design of PermaNet® 3.0, two additional net pieces were cut from the roof panel to provide 7 pieces in total and a representative sample of pyrethroid-PBO incorporated pieces as per WHO recommendation [38]. Net pieces were wrapped in labelled aluminium foil and stored at 30°C before and between use for supplementary cone bioassays and tunnel tests. Following use in laboratory bioassays, net pieces were stored at 4°C before being sent for chemical analysis of insecticide content at the Centre Walloon de Recherches Agronomiques (CRA-W), Belgium.

### Supplementary laboratory bioassays

To provide supplementary data on ITN efficacy, laboratory cone bioassays and tunnel tests were performed with net pieces cut from unwashed and washed ITNs before and after the hut trial. Cone bioassays were performed with the susceptible *An. gambiae s.s.* Kisumu strain to assess the pyrethroid component of ITNs, while tunnel tests were performed with the pyrethroid-resistant *An. gambiae s.l.* Covè strain to assess the chlorfenapyr components of PermaNet® Dual and Interceptor® G2.

- *An. gambiae s.s.* Kisumu strain is an insecticide-susceptible reference strain originated from Kisumu, western Kenya.
- *An. gambiae s.l.* Covè strain are F1 progeny of mosquitoes collected from the experimental hut site in Covè, southern Benin. It is highly resistant to pyrethroids and organochlorines but susceptible to other insecticide classes including chlorfenapyr. Resistance is mediated by the *kdr* L1014F mutation and overexpression of cytochrome P450 monooxygenases [36].

All net pieces cut from unwashed and washed ITNs before and after the hut trial were tested in cone bioassays against the susceptible *An. gambiae s.s.* Kisumu strain. Approximately 10, 2–5-day old mosquitoes were exposed to each net piece for 3 mins in two replicate cones containing ~5 mosquitoes thus giving a total of ~50 mosquitoes per treatment arm. After exposure, mosquitoes were transferred to labelled cups, provided access to 10% (w/v) glucose solution and held at 27±2°C and 75±10% RH. Knockdown was recorded after 60 mins and delayed mortality every 24 h up to 72 h.

Previous studies demonstrate the inability of cone bioassays to predict the field efficacy of chlorfenapyr-based ITNs [39]. To assess the efficacy of the chlorfenapyr component of the pyrethroid-chlorfenapyr nets, we therefore performed tunnel tests against the pyrethroid-resistant Covè with two net pieces randomly selected from those cut from unwashed and washed PermaNet® 2.0, Interceptor® G2 and PermaNet® Dual before and after the hut trial. Tunnel tests are an experimental chamber that mimics the behavioural interactions that occur between free-flying mosquitoes and nets during host-seeking. The design consists of a square glass tunnel divided one third its length by a wooden frame fitted with a net sample. In the short section of the tunnel, a guinea pig bait was held in an open-meshed cage while in the long section, approximately 100, 5–8-day old mosquitoes were released at dusk and left overnight. Net samples were given 9 holes measuring 1 cm in diameter to facilitate entry of mosquitoes into the baited chamber. In the morning, mosquitoes were collected from the tunnel and scored for mortality and blood-feeding. Surviving mosquitoes were placed in labelled plastic cups, provided access to 10% (w/v) glucose solution, and held at 27±2°C and 75±10% RH. Delayed mortality was recorded every 24 h up to 72 h. Similar numbers of mosquitoes were concurrently exposed to untreated net pieces in cone bioassays and tunnel tests as a negative control.

### Chemical analysis of net pieces

Following use in bioassays, all net pieces cut from the selected unwashed and washed ITNs before and after the experimental hut trial were assessed for deltamethrin, alpha-cypermethrin, chlorfenapyr and PBO content.

Deltamethrin and/or chlorfenapyr in PermaNet® Dual, PermaNet® 2.0 and PermaNet® 3.0 (sides) were extracted from net samples by sonication with heptane using dicyclohexyl phthalate as internal standard and determined by normal phase High Performance Liquid Chromatography with UV Diode Array Detection (HPLC-DAD). Alpha-cypermethrin and chlorfenapyr in Interceptor® G2 were extracted from net samples by sonication with heptane using dicyclohexyl phthalate as internal standard and determined by Gas Chromatography with Flame Ionisation Detection (GC-FID).

Deltamethrin in PermaNet® 3.0 (roof) was extracted from net samples by heating under reflux for 30 minutes with xylene using dicyclohexyl phthalate as internal standard. The solvent was evaporated, and the residue dissolved in hexane. Deltamethrin was determined by normal phase High Performance Liquid Chromatography with UV Diode Array Detection (HPLC-DAD). PBO in PermaNet® 3.0 roof was extracted from net samples by heating under reflux for 30 minutes with xylene using octadecane as internal standard and determined by Gas Chromatography with Flame Ionisation Detection (GC-FID).

Each method of analysis was performed using the internal standard calibration. The analytical methods used were based on validated and standardized methods published by the Collaborative International Pesticides Analytical Council (CIPAC). Chemical analysis results were used to calculate proportional retention of active ingredient(s) and synergist after 20 washes.

### Data analysis

Proportional outcomes (mortality, blood-feeding, exophily) were compared between the experimental hut treatments using blocked logistic regression while numerical outcomes (entry) were compared with negative binomial regression. A separate model was fitted for each outcome and adjusted to account for variation between the huts, sleepers and weeks of the trial. In addition, following recent provisional WHO guidance [40], PermaNet® Dual was assessed for its non-inferiority to Interceptor® G2 and its superiority to PermaNet® 2.0 and PermaNet® 3.0 for mosquito mortality and blood-feeding outcomes. Results with unwashed and washed nets were pooled to generate a single efficacy estimate over the lifetime of the net. All analyses were performed in Stata version 17.

### Ethical considerations

Ethical approval for the study was issued by the Research Ethics Committees of the Benin Ministry of Health (Ref: N°34, 09/09/2020) and the London School of Hygiene & Tropical Medicine (LSHTM) (Ref: 26429). Written informed consent was obtained from all human volunteer sleepers prior to participation. Sleepers were offered a free course of chemoprophylaxis spanning the duration of the study and 4 weeks following its completion to mitigate malaria infection risk. Approval for use of guinea pigs for tunnel tests was obtained from the LSHTM Animal Welfare Ethics Review Board (Ref: 2020-01). Guinea pig colonies were maintained at CREC/LSHTM according to standard operating procedures (SOPs) developed in line with relevant national and international regulations governing use of animals for scientific research purposes.

### Compliance with OECD principles of Good Laboratory Practice

To ensure compliance with the OECD principles of GLP, a series of activities were implemented through the initiation, execution, and reporting of the study. The study protocol was developed by a properly trained study director and approved by the sponsor before starting the study. Equipment used for the study (precision balances for weighing insecticides, refrigerators for ITN sample storage and data loggers) were calibrated before use. All ITN products used in the hut trial were verified to be within their expiry dates and were provided with associated certificates of analysis. The candidate net supplied by the manufacturer (Vestergaard Sàrl) was confirmed to come from three production batches. In addition, the environmental conditions under which these products were stored was verified daily by use of a calibrated data logger. Mosquitoes used for cone bioassays and tunnel tests were reared and transported in line with established SOPs that ensured the integrity of the strains tested. All computer systems (data loggers, databases, statistical software) used for data collection, entry, and processing, were validated before use. Records were kept of each procedure performed during the study. The quality assurance team of the CREC/LSHTM Facility performed inspections of the study protocol, critical phases of implementation, data quality and final report to assess compliance to GLP and no non-conformances were detected. The final report, along with all study-related documents, are securely stored in the physical and electronic archive of the Facility for up to 15 years. Study inspections performed in 2021 by the South African National Accreditation System (SANAS), the GLP certification body of the Facility, also detected no non-conformances.

## Results

### WHO bottle bioassay results

Mortality of wild pyrethroid-resistant *An. gambiae s.l.* from the Covè hut station following exposure to the discriminating concentration of deltamethrin was 28% thus confirming the high frequency of pyrethroid resistance in the Covè vector population (Figure 1). Mortality increased progressively with 2x (76%), 5x (96%) and 10x (96%) the discriminating concentration but failed to exceed 98% with any concentration, indicating high intensity deltamethrin resistance. Pre-exposure to PBO fully restored deltamethrin susceptibility (100% mortality) thus suggesting the involvement of cytochrome P450 monooxygenases in pyrethroid resistance. In contrast, chlorfenapyr-treated bottles killed 98% of mosquitoes, indicating full susceptibility to this insecticide. No mortality was recorded in the untreated controls while PBO alone induced 3% mortality.

**Figure 1:**
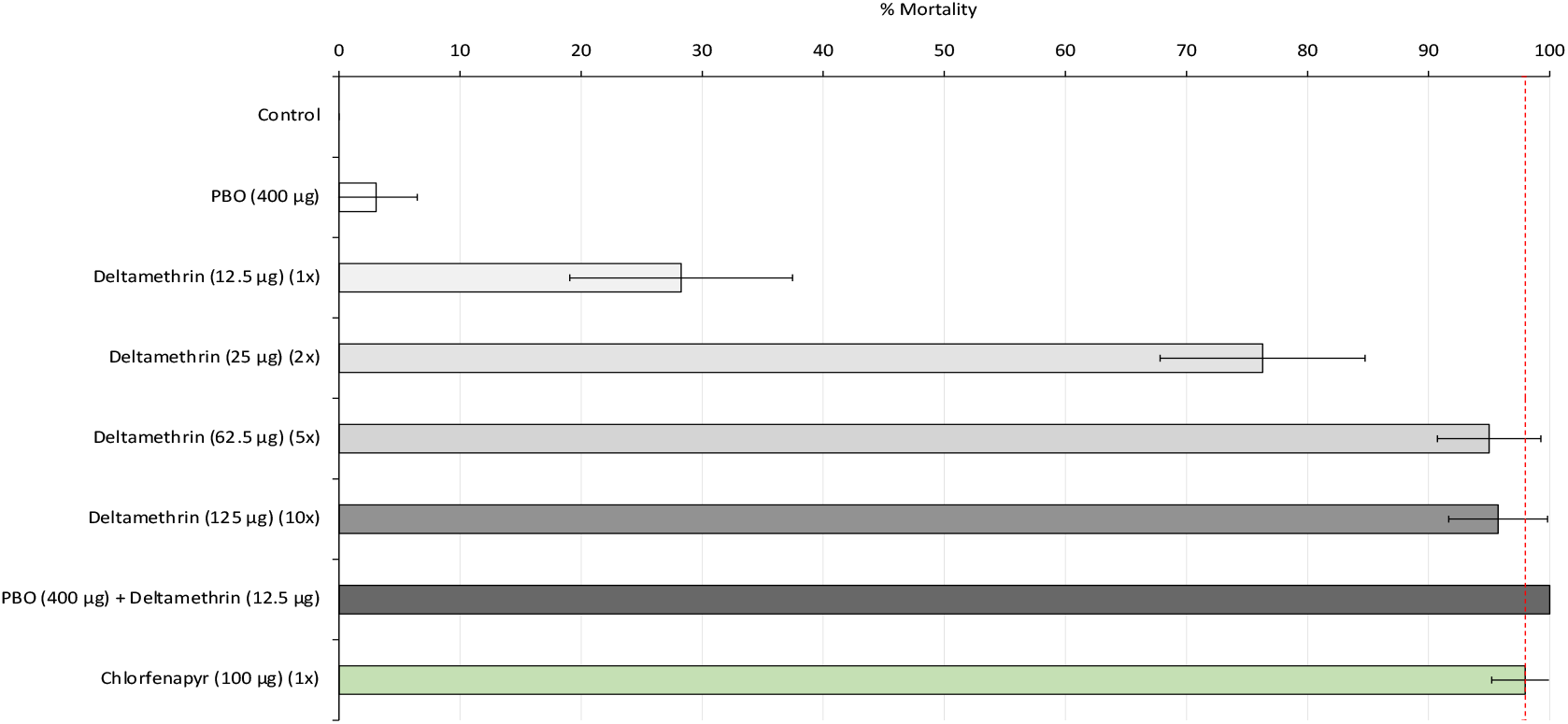
Mortality of F1 progeny of field-collected *Anopheles gambiae sensu lato* in World Health Organisation bottle bioassays. *A total of 80–100 mosquitoes were exposed to each treatment arm for 60 mins in four batches of 20–25. Dashed red line represents 98% susceptibility cut-off and error bars represent 95% CIs.*

### Experimental hut results

#### Entry and exiting results

A total of 5,967 mosquitoes were collected in experimental huts over the 9-week trial, corresponding to an average of approximately 13 mosquitoes per treatment per night (Table 1). None of the ITNs induced a significant deterrent effect relative to the untreated control net and mosquito entry increased significantly with all net types after washing. All ITNs induced significant exiting relative to the control except PermaNet® 2.0 after 20 washes (36% vs. 38%, p=0.584). Exiting was higher with all three dual ITN-types both when unwashed (63%-70%) and after 20 washes (56%-61%) compared to PermaNet® 2.0 (unwashed: 51%, washed: 36%). Mosquito exiting rates did not differ significantly between PermaNet® Dual and Interceptor® G2 or PermaNet® 3.0 both with unwashed nets and nets washed 20 times (p>0.05). Exiting rates generally declined after washing for all net types except Interceptor® G2 (67% vs. 61%, p=0.205).

**Table 1:** Entry and exiting of wild, free-flying, pyrethroid-resistant *Anopheles gambiae sensu lato* entering experimental huts in Covè, southern Benin.

| Net type | Net status | Total females caught* | % Deterrence | Total exiting | % Exophily* | 95% CIs |
| --- | --- | --- | --- | --- | --- | --- |
| Untreated net | — | 541<sup>a</sup> | — | 208 | 38.4<sup>a</sup> | 34.3–42.5 |
| PermaNet® 2.0 | Unwashed | 490<sup>ab</sup> | 9.4 | 248 | 50.6<sup>b</sup> | 46.2–55.0 |
|  | Washed 20x | 903<sup>c</sup> | -66.9 | 329 | 36.4<sup>a</sup> | 33.3–39.6 |
| PermaNet® 3.0 | Unwashed | 591<sup>bd</sup> | -9.2 | 412 | 69.7<sup>c</sup> | 66.0–73.4 |
|  | Washed 20x | 895<sup>c</sup> | -65.4 | 541 | 60.4<sup>d</sup> | 57.2–63.7 |
| Interceptor® G2 | Unwashed | 623<sup>de</sup> | -15.2 | 418 | 67.1<sup>ce</sup> | 63.4–70.8 |
|  | Washed 20x | 669<sup>cd</sup> | -23.7 | 411 | 61.4<sup>def</sup> | 57.7–65.1 |
| PermaNet® Dual | Unwashed | 459<sup>a</sup> | 15.2 | 291 | 63.4<sup>cf</sup> | 59.0–67.8 |
|  | Washed 20x | 796<sup>ce</sup> | -47.1 | 444 | 55.8<sup>d</sup> | 52.3–59.3 |

\*Values in the same column bearing the same letter do not differ significantly at the 5% level according to logistic regression analysis

#### Blood-feeding results

All ITNs significantly reduced blood-feeding relative to the control net except PermaNet® 2.0 washed 20 times (57% vs. 49%, p=0.471) (Figure 2, Table 2). Between unwashed nets, lowest blood-feeding was observed with PermaNet® 3.0 when unwashed (13%) though this increased significantly after washing (30%, p<0.001). Interceptor® G2 induced lower levels of mosquito feeding compared to PermaNet® Dual both before washing (20% vs. 27%, p=0.03) and after washing (32% vs. 39%, p=0.006). Personal protection levels were similar between the pyrethroid-chlorfenapyr nets (52% vs. 54%) when unwashed and declined substantially with both net types after 20 washes albeit to a greater extent with PermaNet® Dual. Nevertheless, PermaNet® Dual provided more blood-feeding inhibition compared to PermaNet® 2.0 before washing (45% vs. 21%) and after 20 washes (21% vs. - 16.5%) (Table 2). For all ITN-types blood-feeding rates were significantly higher with washed nets compared to unwashed nets (p<0.05).

**Figure 2:**
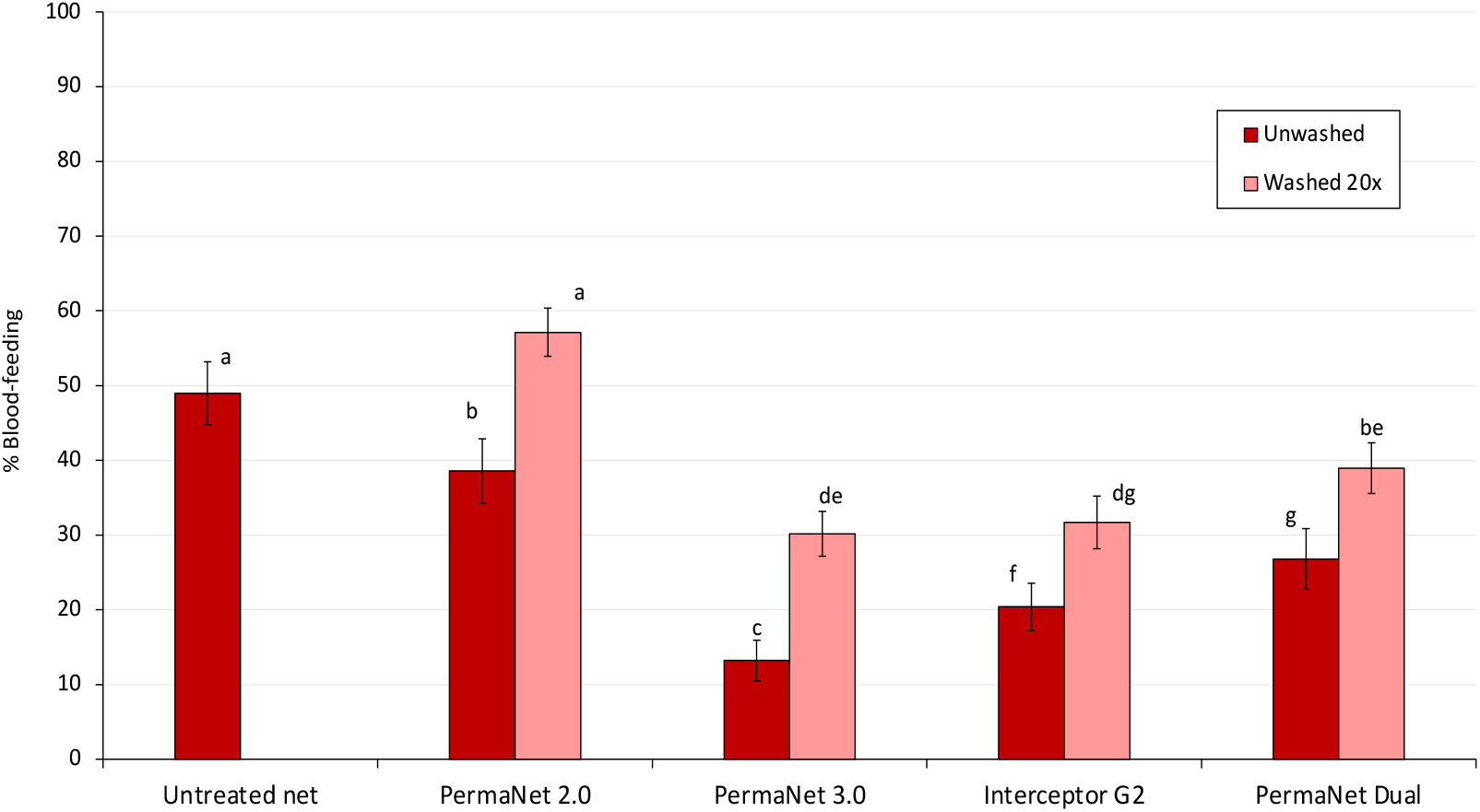
Blood-feeding of wild, free-flying, pyrethroid-resistant *Anopheles gambiae sensu lato* entering experimental huts in Covè, southern Benin. *Bars bearing the same letter do not differ significantly at the 5% level according to logistic regression analysis. Error bars represent 95% CIs.*

**Table 2:** Blood-feeding of wild, free-flying, pyrethroid-resistant *Anopheles gambiae sensu lato* entering experimental huts in Covè, southern Benin.

| Net type | Net status | Total females caught* | Total blood-fed | % Blood-feeding* | 95% CIs | % Blood-feeding inhibition | % Personal protection |
| --- | --- | --- | --- | --- | --- | --- | --- |
| Untreated net | – | 541<sup>a</sup> | 265 | 49.0<sup>a</sup> | 44.8–53.2 | – | – |
| PermaNet® 2.0 | Unwashed | 490<sup>ab</sup> | 189 | 38.6<sup>b</sup> | 34.3–42.9 | 21.3 | 28.7 |
|  | Washed 20x | 903<sup>c</sup> | 516 | 57.1<sup>a</sup> | 53.9–60.4 | -16.7 | -94.7 |
| PermaNet® 3.0 | Unwashed | 591<sup>bd</sup> | 78 | 13.2<sup>c</sup> | 10.5–15.9 | 73.1 | 70.6 |
|  | Washed 20x | 895<sup>c</sup> | 270 | 30.2<sup>de</sup> | 27.2–33.2 | 38.4 | -1.9 |
| Interceptor® G2 | Unwashed | 623<sup>de</sup> | 127 | 20.4<sup>f</sup> | 17.2–23.5 | 58.4 | 52.1 |
|  | Washed 20x | 669<sup>cd</sup> | 212 | 31.7<sup>dg</sup> | 28.2–35.2 | 35.3 | 20.0 |
| PermaNet® Dual | Unwashed | 459<sup>a</sup> | 123 | 26.8<sup>g</sup> | 22.7–30.8 | 45.3 | 53.6 |
|  | Washed 20x | 796<sup>ce</sup> | 310 | 38.9<sup>be</sup> | 35.6–42.3 | 20.5 | -17.0 |

\*Values in the same column bearing the same letter do not differ significantly at the 5% level according to logistic regression analysis

#### Mortality results

Mortality of wild free-flying pyrethroid-resistant *An. gambiae s.l.* with the untreated control net was 2% (Figure 3, Table 3). Among the ITNs, lowest mosquito mortality was achieved with PermaNet® 2.0 (unwashed: 23%, washed: 14%). PermaNet® 3.0 induced higher mortality than PermaNet® 2.0 both with unwashed nets (56% vs. 23%, p<0.001) and nets washed 20 times (30% vs. 14%, p<0.001). Mortality decreased significantly after washing with both PermaNet® 2.0 (23% vs. 14%, p=0.002) and PermaNet® 3.0 (56% vs. 30%, p<0.001). The pyrethroid-chlorfenapyr nets induced significantly higher levels of mosquito mortality (76%-83% when unwashed and 75% with both net types after 20 washes) compared to PermaNet® 2.0 and PermaNet® 3.0 (p<0.001). Interceptor® G2 induced higher vector mortality than PermaNet® Dual when unwashed (83% vs. 76%, p=0.019) but similar mortality after 20 washes (75% vs. 75%, p=0.865). While a significant decline in vector mortality was observed with Interceptor® G2 after 20 washes (83% to 75%, p=0.002), the levels of mortality achieved with PermaNet® Dual remained the same after washing (76% vs. 75%, p=0.684).

**Figure 3:**
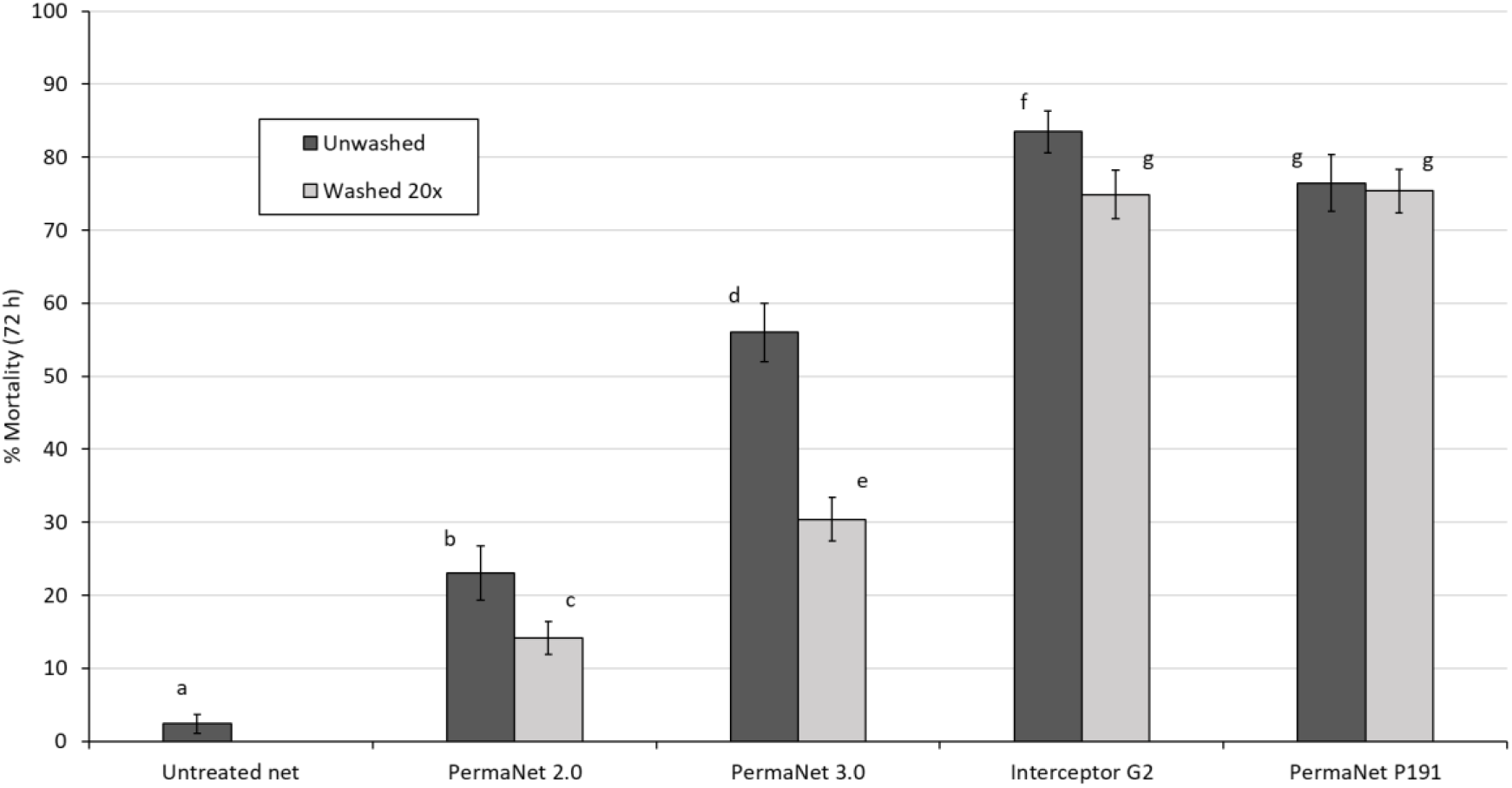
Mortality (72 h) of wild, free-flying, pyrethroid-resistant *Anopheles gambiae sensu lato* entering experimental huts in Covè, southern Benin. *Bars bearing the same letter do not differ significantly at the 5% level according to logistic regression analysis. Error bars represent 95% CIs.*

**Table 3:**
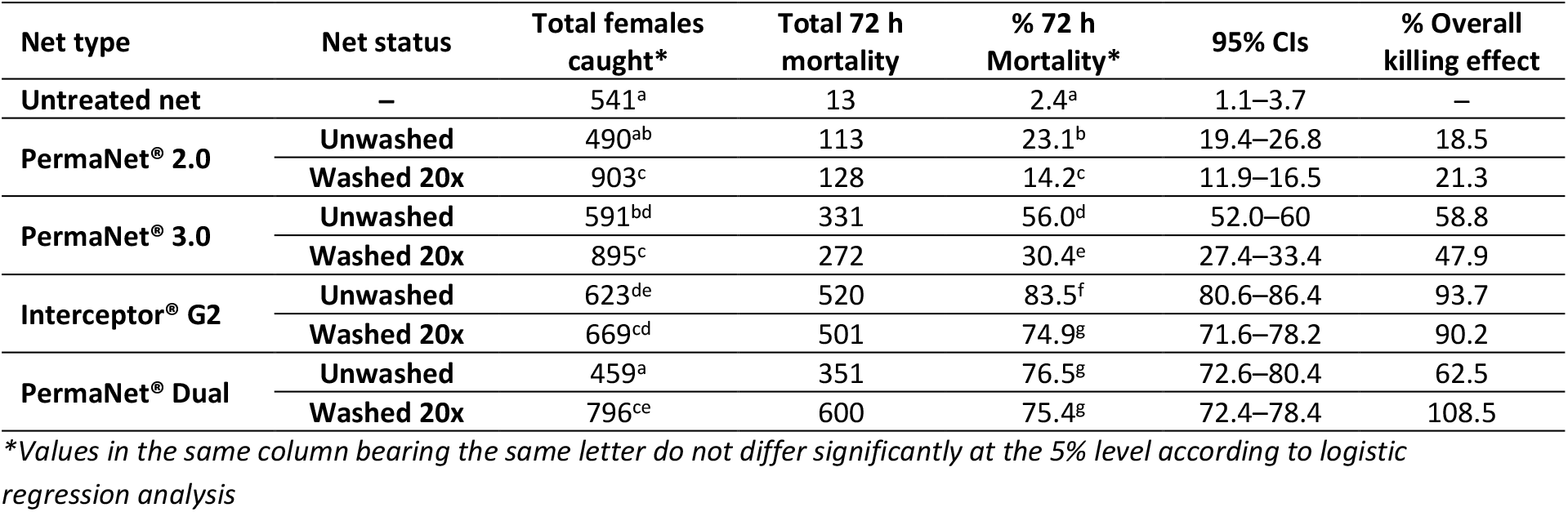
Mortality of wild, free-flying, pyrethroid-resistant *Anopheles gambiae* sensu lato entering experimental huts in Covè, southern Benin.

#### Non-inferiority assessment

Following provisional WHO guidelines recommending a non-inferiority margin of 0.7 [40], PermaNet® Dual was considered non-inferior to Interceptor® G2 for mortality if the lower 95% confidence interval (CI) of the odds ratio describing the difference in mortality was greater than 0.7 and for blood-feeding if the upper 95% CI estimate of the odds ratio describing the difference in blood-feeding was lower than 1.43. PermaNet® Dual was also tested for its superiority over PermaNet® 2.0 and PermaNet® 3.0 for mortality and blood-feeding outcomes. For the non-inferiority and superiority assessments, results with unwashed and washed nets were pooled to generate a single efficacy estimate over the lifetime of the net. As per the recommendations of a recent WHO technical consultation [41], mortality was adopted as the primary endpoint to assess the non-inferiority of PermaNet® Dual while blood-feeding was included as a secondary endpoint to support programmatic decision-making.

The odds ratio describing the difference between PermaNet® Dual and Interceptor® G2 was 0.854 for mortality (76% vs. 79%, 95% CIs: 0.703–1.038) and 1.445 for blood-feeding (35% vs. 26%, 95% CIs: 1.203–1.735) (Figures 4 & 5, Table 5). Based on the non-inferiority margin outlined above, PermaNet® Dual was therefore non-inferior to Interceptor® G2 in terms of its ability to kill vector mosquitoes but not non-inferior in terms of its ability to prevent blood-feeding. PermaNet® Dual demonstrated superiority over PermaNet® 2.0 both in terms of vector mortality (76% vs. 17%, p<0.001) and blood-feeding (35% vs. 51%, p<0.001). PermaNet® Dual was also superior to PermaNet® 3.0 in terms of inducing vector mortality (76% vs. 41%, p<0.001) however, it was inferior in terms of preventing blood-feeding (35% vs. 23%, p<0.001). Detailed results from the non-inferiority and superiority assessments are provided in Table S1.

**Figure 4:**
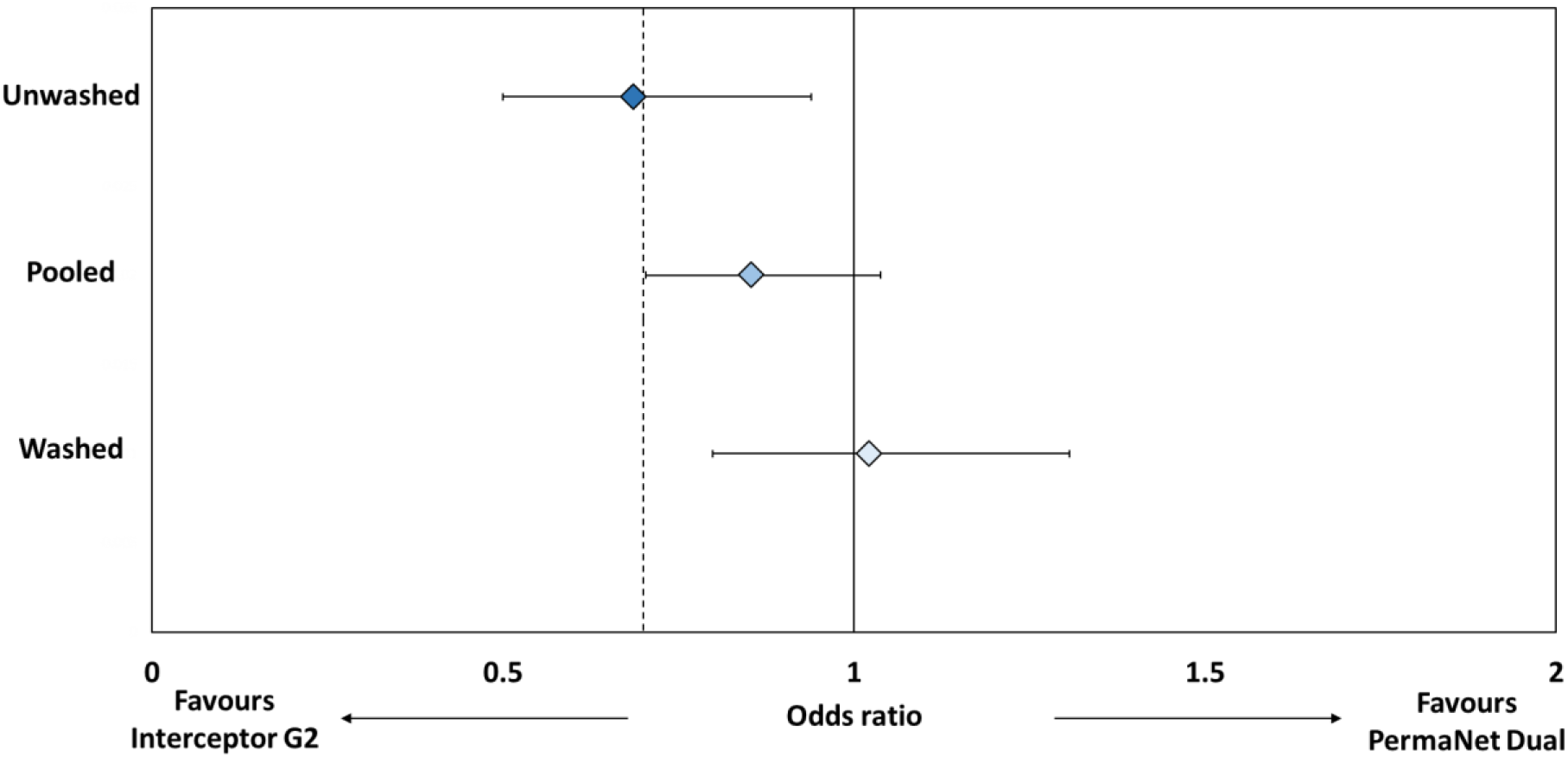
Non-inferiority analysis for proportional mosquito mortality in experimental huts for PermaNet® Dual compared to Interceptor® G2. *Odds ratios represented by blue-shaded diamonds for unwashed nets, washed nets and pooled analysis. Error bars represent 95% CIs. Dashed line represents margin of non-inferiority (odds ratio=0.7). Lower 95% CI must exceed dashed line to fulfill non-inferiorty criteria.*

**Figure 5:**
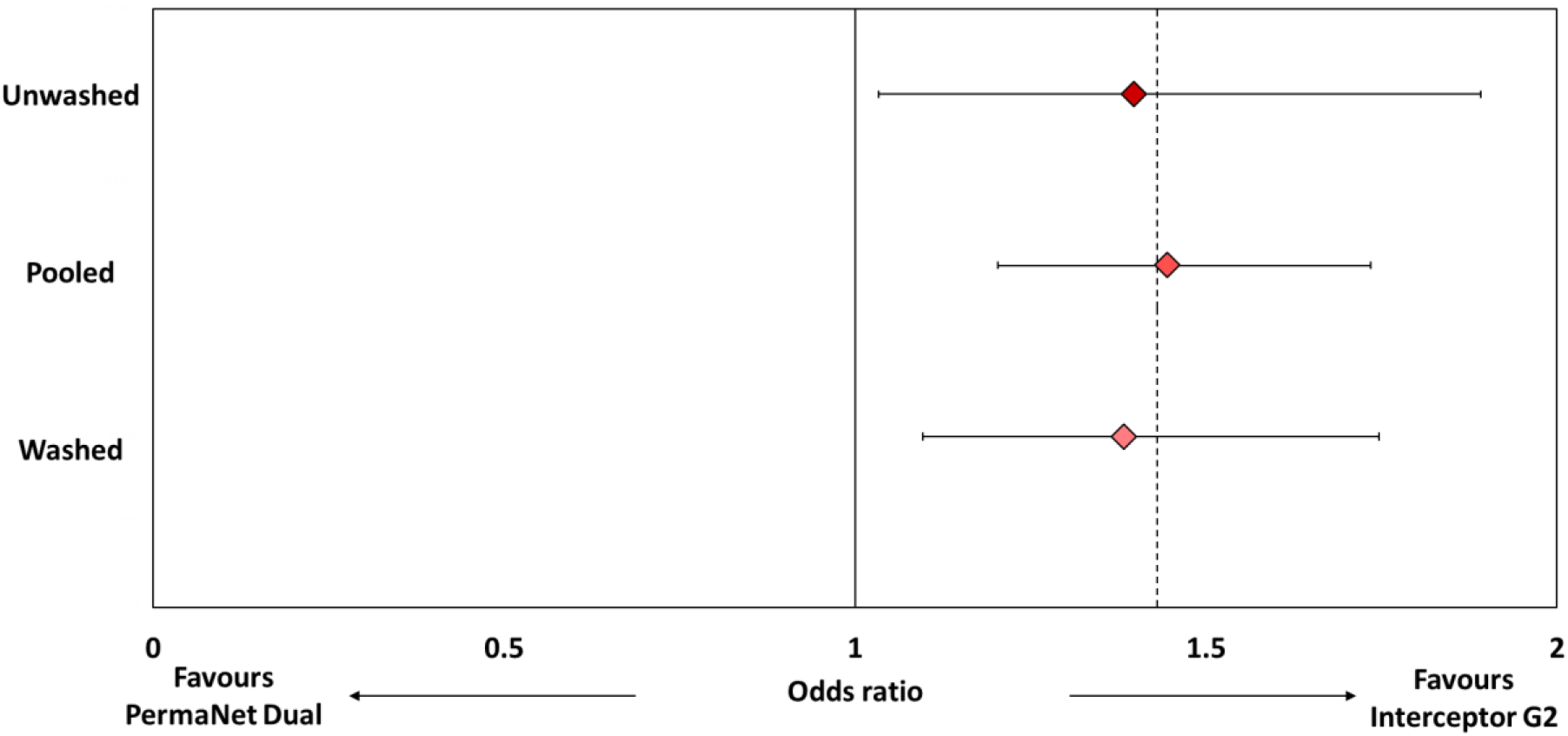
Non-inferiority analysis for proportional mosquito blood-feeding in experimental huts for PermaNet® Dual compared to Interceptor® G2. *Odds ratios represented by red-shaded diamonds for unwashed nets, washed nets and pooled analysis. Error bars represent 95% CIs. Dashed line represents margin of non-inferiority (odds ratio=1.43). Upper 95% CI must not exceed dashed line to fulfill non-inferiorty criteria.*

**Table 5:** Non-inferiority analyses comparing the effect of PermaNet® Dual to Interceptor® G2 for mosquito mortality and blood-feeding outcomes in experimental huts. *To fulfill non-inferiority criteria, lower 95% CI of odds ratio must exceed 0.7 for mortality while upper 95% CI of odds ratio must not exceed 1.43 for blood-feeding.*

|  | Unwashed |  | Washed |  | Pooled |  |
| --- | --- | --- | --- | --- | --- | --- |
|  | Odds ratio<br>(95% CIs) | Non-inferiority | Odds ratio<br>(95% CIs) | Non-inferiority | Odds ratio<br>(95% CIs) | Non-inferiority |
| **Mortality** | 0.686<br>(0.500–0.939) | Not non-inferior | 1.022<br>(0.799–1.307) | Non-inferior | 0.854<br>(0.703–1.038) | Non-inferior |
| **Blood-feeding** | 1.398<br>(1.034–1.891) | Not non-inferior | 1.383<br>(1.096–1.746) | Not non-inferior | 1.445<br>(1.203–1.735) | Not non-inferior |

### Supplementary laboratory bioassay results

Cone bioassay results with the susceptible *An. gambiae s.s.* Kisumu strain are provided in Figure 5 with more detailed results provided in supplementary information (Table S2). PermaNet® 3.0 roof samples induced the highest mortality rates in cone bioassays (78–97%). As expected, the performance of the pyrethroid-chlorfenapyr nets was very poor in cone bioassays inducing <60% mortality with all ITN pieces tested. Hence, the results further demonstrate the unsuitability of cone bioassays for testing pyrethroid-chlorfenapyr ITNs.

Tunnel test mortality with the pyrethroid-resistant Covè strain was lowest with PermaNet® 2.0, (<70%) though this did not decline significantly after washing (Figure 6). In contrast, unwashed and washed net pieces of both pyrethroid-chlorfenapyr ITNs taken before and after the hut trial induced ≥98% mortality. Blood-feeding inhibition was high with all ITNs (50–93%). Highest blood-feeding inhibition was achieved with PermaNet® Dual, exceeding 85% with all net pieces and was similar between unwashed and washed pieces taken before (88% vs. 91%) and after the hut trial (93% vs. 90%). More detailed results from the tunnel tests are provided in supplementary information (Table S3).

**Figure 6:**
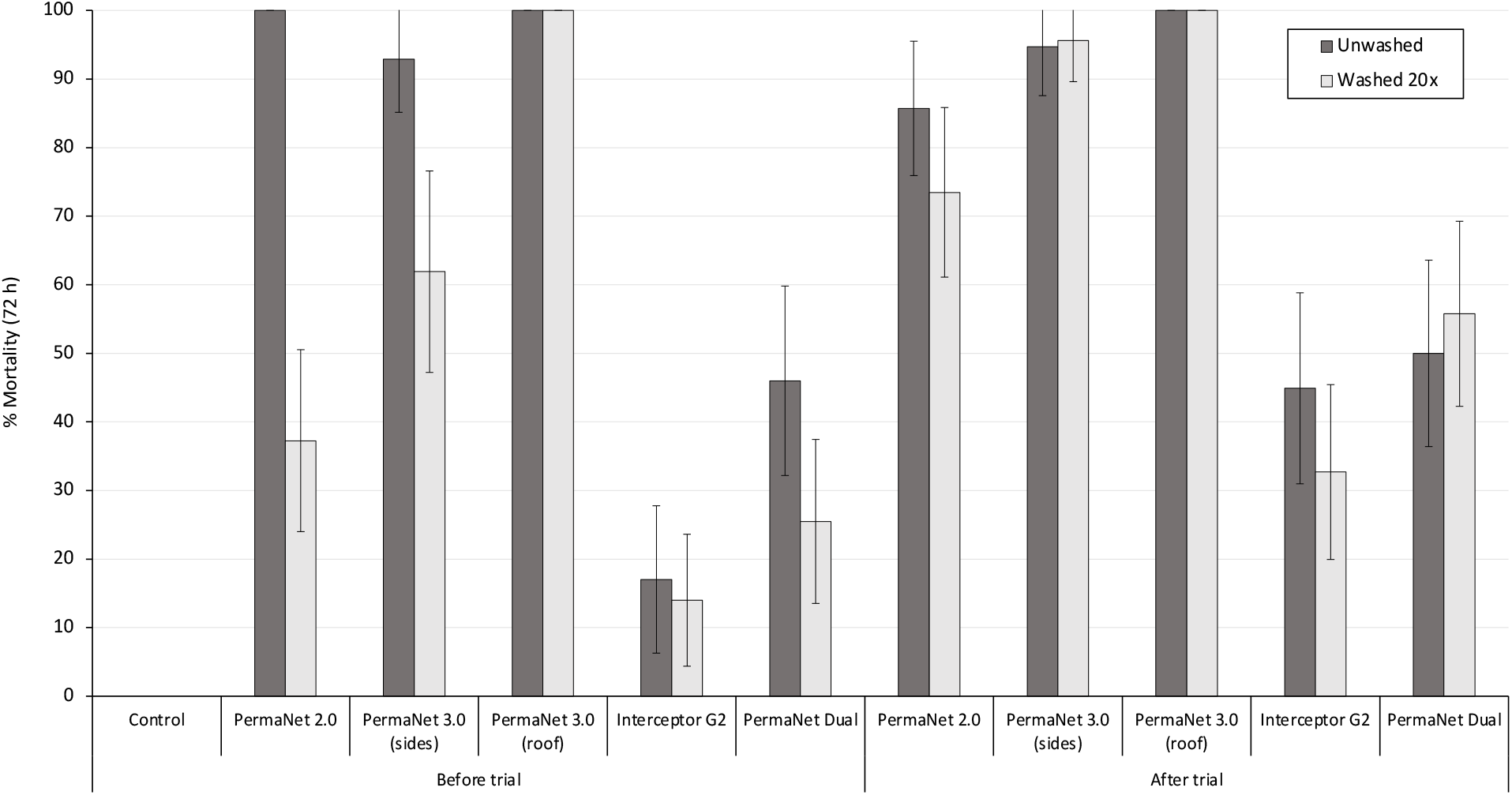
Mortality after 72 h of susceptible *Anopheles gambiae sensu stricto* Kisumu strain in supplementary cone bioassays. *Approximately 10 mosquitoes were exposed to each of the 5 net pieces cut from unwashed and washed nets before and after the hut trial for 3 mins in two batches of 5. Error bars represent 95% CIs.*

**Figure 7:**
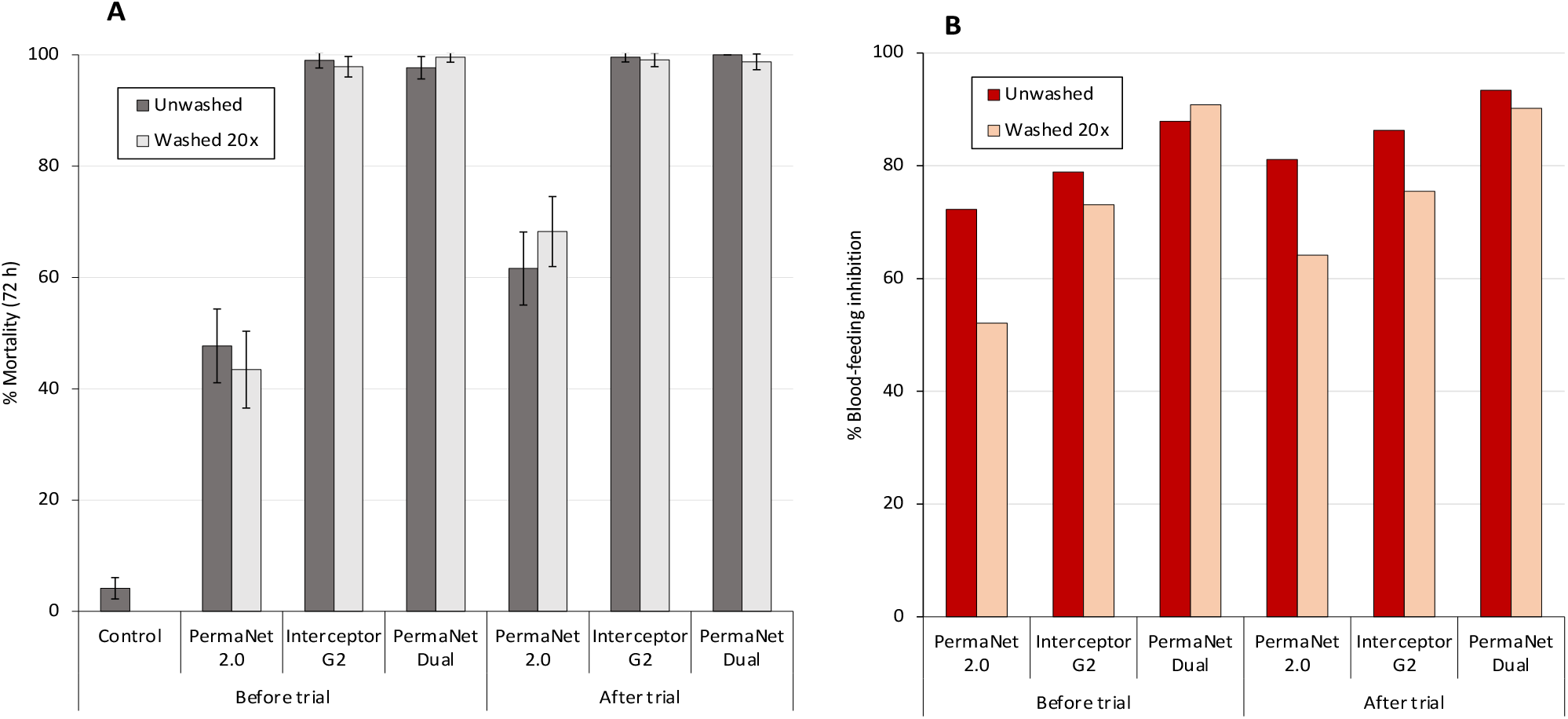
Mortality after 72 h (A) and blood-feeding inhibition (B) of pyrethroid-resistant *Anopheles gambiae sensu lato* Covè strain in supplementary tunnel tests. *Approximately 100 mosquitoes were exposed to each of the two randomly selected net pieces from each treatment arm overnight in one replicate tunnel test. Error bars represent 95% CIs.*

### Chemical analysis of net pieces results

The active ingredient content in all unwashed ITNs were within defined specifications declared by the manufacturers. Retention of deltamethrin after 20 washes was lowest with net pieces cut from PermaNet® 2.0 (17%) (Table 6). PermaNet® 3.0 showed higher proportional wash-retention of deltamethrin on the roof panel (86.1%) compared to the side panels (26%). The PBO component showed moderate levels of wash-retention (60%). Between the pyrethroid-chlorfenapyr ITNs, Interceptor® G2 showed higher levels of wash-retention of both active ingredients (87% for alpha-cypermethrin and 65% for chlorfenapyr) compared to PermaNet® Dual (42% for deltamethrin and 25% for chlorfenapyr). The wash-resistance index of chlorfenapyr was thus higher with Interceptor® G2 (97.9%) than with PermaNet® Dual (93.3%).

**Table 6:**
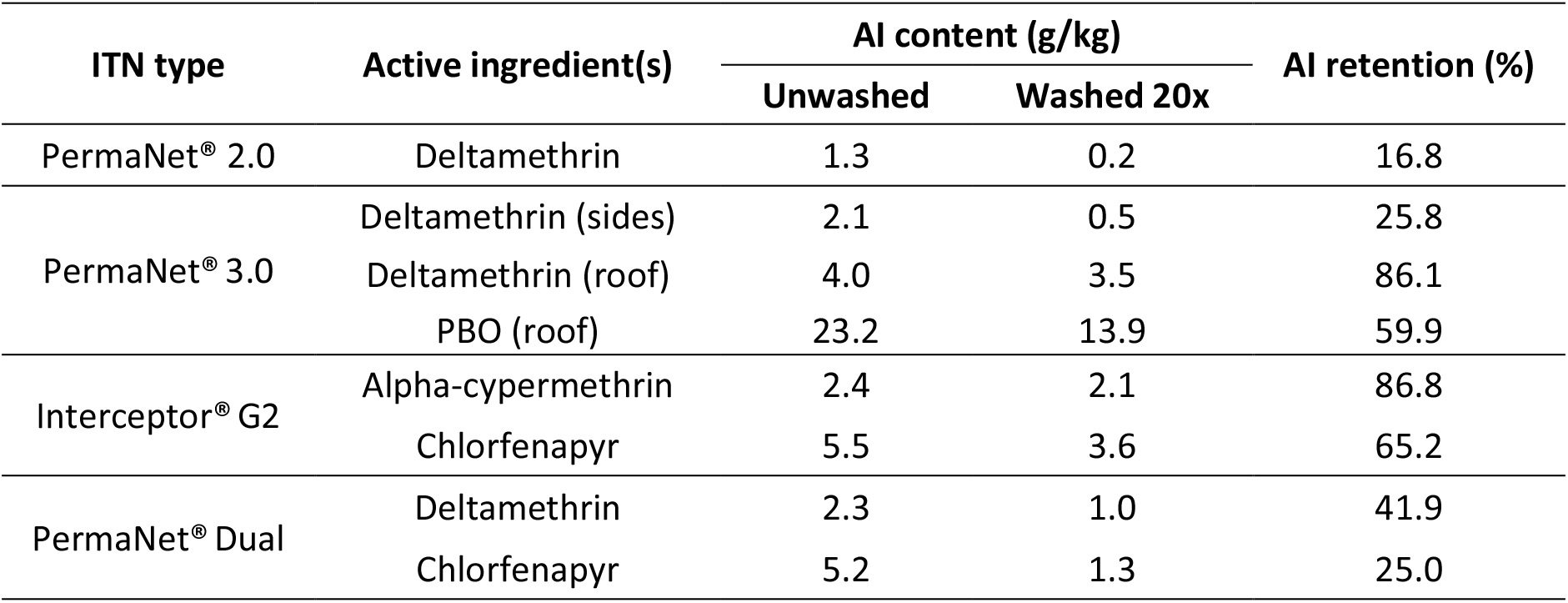
Chemical content of unwashed and washed net pieces taken before and after the experimental hut trial in Covè, Benin.

## Discussion

This study evaluated the efficacy and wash resistance of PermaNet® Dual – a new deltamethrin-chlorfenapyr net – against a pyrethroid-resistant malaria vector population in an experimental hut trial in southern Benin. PermaNet® Dual was investigated for its superiority to WHO-prequalified pyrethroid-only (PermaNet® 2.0) and pyrethroid-PBO (PermaNet® 3.0) ITNs and non-inferiority to a WHO-prequalified pyrethroid-chlorfenapyr ITN (Interceptor® G2) with empirical evidence of public health value.

The poor performance of PermaNet® 2.0 (<25% mortality) is typical of experimental hut trials conducted with pyrethroid-only nets at the Covè hut site and is attributable to the high intensity of pyrethroid resistance demonstrated in susceptibility bottle bioassays in this study and in previous [35, 36]. Complete restoration of susceptibility to deltamethrin following pre-exposure to PBO was observed in the bottle bioassays which suggests strong involvement of cytochrome P450 monooxygenase activity in deltamethrin resistance in the Covè vector population during the hut trial. Previous bioassays performed with wild *An. gambiae s.l.* from Covè using different pyrethroid insecticides have usually resulted in partial or no restoration of susceptibility to pyrethroids with pre-exposure to PBO [42]. This variability in outcome of synergist bioassays may be due to differences in the type of pyrethroid insecticide tested, test methods or seasonal changes in the vector population. However, despite complete restoration of pyrethroid susceptibility following PBO pre-exposure in bottle bioassays in this hut trial, the levels of improved mosquito mortality achieved with the PermaNet® 3.0 relative to PermaNet® 2.0 were moderate (17% vs. 40%) and did not differ substantially compared to what has been reported in previous hut studies with this vector population [20, 43]. This finding would suggest the presence of more complex behavioural mechanisms that may have reduced mosquito contact with PermaNet® 3.0 compromising its efficacy in the experimental huts. Further studies to investigate the relationship between levels of restoration of susceptibility to pyrethroids achieved in PBO pre-exposure bioassays and the efficacy of pyrethroid-PBO ITNs would be useful.

Both pyrethroid-chlorfenapyr ITNs (PermaNet® Dual and Interceptor® G2) induced significantly higher levels of mortality (75%-86%) of wild pyrethroid-resistant malaria vector mosquitoes entering the experimental huts relative to the pyrethroid-only and pyrethroid-PBO ITNs. This is mostly due to susceptibility of the Covè vector population to chlorfenapyr as demonstrated in bottle bioassays. This superior performance of pyrethroid-chlorfenapyr ITNs confirms previous findings in experimental hut studies in Benin [22, 44] and across Africa [23–25] and recent cRCTs in Benin [27] Tanzania [28], reiterating the importance of this innovative ITN technology for improving the control of pyrethroid-resistant malaria vector populations. PermaNet® Dual was also non-inferior to Interceptor® G2 for the primary endpoint of mortality thus providing necessary evidence for the candidate ITN to be covered by WHO policy recommendations for deployment of pyrethroid-chlorfenapyr nets, pending their availability. Studies investigating its entomological performance against other malaria vector species in other ecological settings are ongoing and will add to the body of evidence to support its deployment. Prequalification of PermaNet® Dual by WHO will provide additional choice of pyrethroid-chlorfenapyr nets to vector control programmes and help procurers meet increasing global demand for this effective dual-active ITN class by endemic countries.

Experimental hut performance after 20 washes is used as a proxy for ITN efficacy after 3 years of field use [37]. Although wash-retention of chlorfenapyr was lower in PermaNet® Dual relative to Interceptor® G2, its performance in experimental huts remained unchanged after 20 standardised washes, showing potential for the net to demonstrate durable bioefficacy. This finding was supported by the tunnel tests demonstrating high mortality of pyrethroid-resistant Covè mosquitoes (>95%) with the two pyrethroid-chlorfenapyr ITNs, both before and after 20 washes. However, further studies to monitor the post-market performance of PermaNet® Dual including assessment of its fabric integrity, bioefficacy and chemical content under household use over 3 years, are advisable.

While a superior performance of pyrethroid-chlorfenapyr nets relative to pyrethroid-only and pyrethroid-PBO nets was clearly demonstrated in this study and in previous hut studies and cRCTs, care should be taken not to over-rely on this one class of insecticide as this may quickly drive development of resistance to chlorfenapyr eventually leading to product failure. Pyrethroid-chlorfenapyr nets should be ideally deployed alongside other insecticide chemistries or in rotation with other ITN types as part of an insecticide resistance management strategy aimed at preventing the selection of chlorfenapyr resistance and extending the useful life of this ITN class.

## Conclusions

PermaNet® Dual, a new deltamethrin-chlorfenapyr ITN developed by Vestergaard Sàrl, demonstrated superior performance compared to a pyrethroid-only ITN (PermaNet® 2.0) and a pyrethroid-PBO (PermaNet® 3.0) ITN in experimental huts against wild, free-flying pyrethroid-resistant *An gambiae s.l.* in Benin. PermaNet® Dual was also non-inferior to Interceptor® G2, a WHO-prequalified pyrethroid-chlorfenapyr ITN that has demonstrated evidence of improved public health impact in cluster randomised-controlled trials. The addition of PermaNet® Dual to the current WHO list of prequalified ITNs presents an additional option of this highly effective ITN class for improved control of malaria transmitted by pyrethroid-resistant mosquito vectors.

## List of abbreviations

PBO: Piperonyl butoxide
ITN: Insecticide treated nets
LLIN: Long-lasting insecticidal nets
WHO: World Health Organization
PQ: Prequalification team
cRCT: Cluster randomised controlled trial
GLP: Good laboratory practice
CREC: Centre de Recherche Entomologique de Cotonou
LSHTM: London School of Hygiene & Tropical Medicine

## Availability of data and material

The datasets used and/or analysed during the current study are available from the corresponding author on reasonable request.

## Authors’ contributions

CN designed the study, acquired funding, supervised the project and prepared the final manuscript. TS supervised the hut trial, analysed the data, prepared the graphs and contributed to manuscript preparation. BN and MG performed the hut trial and laboratory bioassays. DT performed the susceptibility bioassays. VA ensured compliance to principles of principles of Good Laboratory Practice. OP and PDV performed the chemical analysis. All authors read and approved the final version of the manuscript.

## Acknowledgements

We thank Dr. Eleanore Sternberg (previously Vestergaard Sàrl) and Melinda Hadi of Vestergaard Sàrl for providing the PermaNet® nets. We also thank the staff of CREC/LSHTM (Renaud Govoetachan, Josias Fagbohoun, Estelle Vigninou, Abel Agbevo etc.) for their assistance and Ms. Imelda Glele and Danielle Apithy for administrative and logistics support. Special thanks to the rice farmers at Covè for their participation in the hut study.

## References

1. Pryce, J., M. Richardson, and C. Lengeler, Insecticide-treated nets for preventing malaria. Cochrane Database of Systematic Reviews, 2018 (11).

2. Lim, S.S., et al., Net benefits: a multicountry analysis of observational data examining associations between insecticide-treated mosquito nets and health outcomes. PLoS medicine, 2011. 8(9): p. e1001091.

3. Bhatt, S., et al., The effect of malaria control on Plasmodium falciparum in Africa between 2000 and 2015. Nature, 2015. 526(7572): p. 207–211.

4. World Health Organisation, World malaria report 2021. 2021, World Health Organisation.

5. Kleinschmidt, I., et al., Implications of insecticide resistance for malaria vector control with long-lasting insecticidal nets: a WHO-coordinated, prospective, international, observational cohort study. The Lancet infectious diseases, 2018. 18(6): p. 640–649.

6. Strode, C., et al., The impact of pyrethroid resistance on the efficacy of insecticide-treated bed nets against African anopheline mosquitoes: systematic review and meta-analysis. PLoS Med, 2014. 11(3): p. e1001619.

7. Toé, K.H., et al., Increased pyrethroid resistance in malaria vectors and decreased bed net effectiveness, Burkina Faso. Emerging infectious diseases, 2014. 20(10): p. 1691.

8. Asidi, A., et al., Loss of household protection from use of insecticide-treated nets against pyrethroid-resistant mosquitoes, Benin. Emerging infectious diseases, 2012. 18(7): p. 1101.

9. N’Guessan, R., et al., Reduced efficacy of insecticide-treated nets and indoor residual spraying for malaria control in pyrethroid resistance area, Benin. Emerging infectious diseases, 2007. 13(2): p. 199.

10. Bingham, G., et al., Can piperonyl butoxide enhance the efficacy of pyrethroids against pyrethroid-resistant Aedes aegypti? Tropical Medicine & International Health, 2011. 16(4): p. 492–500.

11. Oumbouke, W.A., et al., Evaluation of an alpha-cypermethrin+ PBO mixture long-lasting insecticidal net VEERALIN® LN against pyrethroid resistant Anopheles gambiae ss: an experimental hut trial in M’bé, central Côte d’Ivoire. Parasites & vectors, 2019. 12(1): p. 1–10.

12. Akoton, R., et al., Experimental huts trial of the efficacy of pyrethroids/piperonyl butoxide (PBO) net treatments for controlling multi-resistant populations of Anopheles funestus ss in Kpomè, Southern Benin. Wellcome Open Research, 2018. 3.

13. Toe, K., et al., Do bednets including piperonyl butoxide offer additional protection against populations of Anopheles gambiae sl. that are highly resistant to pyrethroids? An experimental hut evaluation in Burkina Fasov. Medical and veterinary entomology, 2018. 32(4): p. 407–416.

14. N’Guessan, R., et al., An experimental hut evaluation of PermaNet® 3.0, a deltamethrin— piperonyl butoxide combination net, against pyrethroid-resistant Anopheles gambiae and Culex quinquefasciatus mosquitoes in southern Benin. Transactions of the Royal Society of Tropical Medicine and Hygiene, 2010. 104(12): p. 758–765.

15. Tungu, P., et al., Evaluation of PermaNet 3.0 a deltamethrin-PBO combination net against Anopheles gambiae and pyrethroid resistant Culex quinquefasciatus mosquitoes: an experimental hut trial in Tanzania. Malaria Journal, 2010. 9(1): p. 1–13.

16. Protopopoff, N., et al., Effectiveness of a long-lasting piperonyl butoxide-treated insecticidal net and indoor residual spray interventions, separately and together, against malaria transmitted by pyrethroid-resistant mosquitoes: a cluster, randomised controlled, two-by-two factorial design trial. Lancet, 2018. 391(10130): p. 1577–1588.

17. Staedke, S.G., et al., Effect of long-lasting insecticidal nets with and without piperonyl butoxide on malaria indicators in Uganda (LLINEUP): a pragmatic, cluster-randomised trial embedded in a national LLIN distribution campaign. Lancet, 2020. 395(10232): p. 1292–1303.

18. World Health Organisation, Guidelines for malaria. 2022, World Health Organisation.

19. Mechan, F., et al., LLIN Evaluation in Uganda Project (LLINEUP)–The durability of long-lasting insecticidal nets treated with and without piperonyl butoxide (PBO) in Uganda. bioRxiv, 2022.

20. Syme, T., et al., Pyrethroid-piperonyl butoxide (PBO) nets reduce the efficacy of indoor residual spraying with pirimiphos-methyl against pyrethroid-resistant malaria vectors. Sci Rep, 2022. 12(1): p. 6857.

21. World Health Organisation, List of WHO prequalified vector control products. 2023, World Health Organisation

22. N’Guessan, R., et al., A chlorfenapyr mixture net interceptor® G2 shows high efficacy and wash durability against resistant mosquitoes in West Africa. PLoS One, 2016. 11(11): p. e0165925.

23. Bayili, K., et al., Evaluation of efficacy of Interceptor® G2, a long-lasting insecticide net coated with a mixture of chlorfenapyr and alpha-cypermethrin, against pyrethroid resistant Anopheles gambiae sl in Burkina Faso. Malaria journal, 2017. 16(1): p. 1–9.

24. Camara, S., et al., Efficacy of Interceptor® G2, a new long-lasting insecticidal net against wild pyrethroid-resistant Anopheles gambiae ss from Côte d’Ivoire: a semi-field trial. Parasite, 2018. 25.

25. Tungu, P.K., et al., Efficacy of interceptor® G2, a long-lasting insecticide mixture net treated with chlorfenapyr and alpha-cypermethrin against Anopheles funestus: experimental hut trials in north-eastern Tanzania. Malaria Journal, 2021. 20(1): p. 1–15.

26. Oxborough, R.M., et al., ITN mixtures of chlorfenapyr (pyrrole) and alphacypermethrin (pyrethroid) for control of pyrethroid resistant Anopheles arabiensis and Culex quinquefasciatus. PLoS One, 2013. 8(2): p. e55781.

27. Accrombessi, M., et al., Efficacy of pyriproxyfen-pyrethroid long-lasting insecticidal nets (LLINs) and chlorfenapyr-pyrethroid LLINs compared with pyrethroid-only LLINs for malaria control in Benin: a cluster-randomised, superiority trial. The Lancet, 2023.

28. Mosha, J.F., et al., Effectiveness and cost-effectiveness against malaria of three types of dual-active-ingredient long-lasting insecticidal nets (LLINs) compared with pyrethroid-only LLINs in Tanzania: a four-arm, cluster-randomised trial. The Lancet, 2022. 399(10331): p. 1227–1241.

29. Accrombessi, M., et al., Assessing the efficacy of two dual-active ingredients long-lasting insecticidal nets for the control of malaria transmitted by pyrethroid-resistant vectors in Benin: study protocol for a three-arm, single-blinded, parallel, cluster-randomized controlled trial. BMC Infect Dis, 2021. 21(1): p. 194.

30. AMP, Alliance for malaria prevention; mass campagn tracker. https://allianceformalariaprevention.com/mass-campaign-tracker/?_sfm_mc_date_of_import=20221015, 2022. Accessed 28th October 2022.

31. UNICEF, Long-lasting insecticidal nets market update. https://www.unicef.org/supply/media/13951/file/LLIN-Market-and-Supply-Update-October-2022.pdf, 2020. accessed 27th October 2022.

32. WHO, Data requirements and protocol for determining non-inferiority of insecticide-treated net and indoor residual spraying products within an established WHO intervention class. Geneva, Switzerland, 2018. https://apps.who.int/iris/bitstream/handle/10665/276039/WHO-CDS-GMP-2018.22-eng.pdf?ua=1.

33. WHO, Norms, standards and processes underpinning development of WHO recommendations on vector control 22 December 2020. https://www.who.int/publications/i/item/9789240017382, 2020.

34. WHO, Standard operating procedure for testing insecticide susceptibility of adult mosquitoes in WHO bottle bioassays, WHO Bottle-bioassay/01/14 January 2022. World Health Organisation, Geneva, 2022. https://apps.who.int/iris/handle/10665/352312?search-result=true&query=WHO+bottie+bioassav&scope=&rpp=10&sort_by=score&order=desc.

35. Syme, T., et al., Which indoor residual spraying insecticide best complements standard pyrethroid long-lasting insecticidal nets for improved control of pyrethroid resistant malaria vectors? PloS one, 2021. 16(1): p. e0245804.

36. Ngufor, C., et al., Insecticide resistance profile of Anopheles gambiae from a phase II field station in Cove, southern Benin: implications for the evaluation of novel vector control products. Malar J, 2015. 14: p. 464.

37. World Health Organisation, Guidelines for laboratory and field-testing of long-lasting insecticidal nets. 2013, World Health Organisation.

38. World Health Organisation, WHO Specifications and Evaluations for Public Health Pesticides Deltamethrin + Piperonyl Butoxide Long-Lasting (Incorporated Into Filaments) Insecticidal Net. 2012, World Health Organisation.

39. Oxborough, R.M., et al., The activity of the pyrrole insecticide chlorfenapyr in mosquito bioassay: towards a more rational testing and screening of non-neurotoxic insecticides for malaria vector control. Malaria journal, 2015. 14(1): p. 1–11.

40. World Health Organisation, Data requirements and protocol for determining non-inferiority of insecticide-treated net and indoor residual spraying products within an established WHO intervention class. 2018, World Health Organisation.

41. World Health Organisation, Technical consultation on determining non-inferiority of vector control products within an estbalished class. 2021, World Health Organisation.

42. Mosha, J.F., et al., Effectiveness and cost-effectiveness against malaria of three types of dual-active-ingredient long-lasting insecticidal nets (LLINs) compared with pyrethroid-only LLINs in Tanzania: a four-arm, cluster-randomised trial. Lancet, 2022. 399(10331): p. 1227–1241.

43. Ngufor, C., et al., Comparative efficacy of two pyrethroid-piperonyl butoxide nets (Olyset Plus and PermaNet 3.0) against pyrethroid resistant malaria vectors: a non-inferiority assessment. Malar J, 2022. 21(1): p. 20.

44. Ngufor, C., et al., Which intervention is better for malaria vector control: insecticide mixture long-lasting insecticidal nets or standard pyrethroid nets combined with indoor residual spraying? Malar J, 2017. 16(1): p. 340.

