## Supplemental Tables 1-3 for "PermaNet® Dual, a new deltamethrin-chlorfenapyr mixture net, shows improved efficacy against pyrethroid-resistant *Anopheles gambiae sensu lato* in southern Benin"

**Table S1:** Non-inferiority and superiority analyses comparing the effect of PermaNet® Dual to Interceptor® G2, PermaNet® 2.0 and PermaNet® 3.0 for mosquito mortality and blood-feeding outcomes in experimental huts.

| | | Superiority assessments | | | | Non-inferiority assessment | |
|---|---|---|---|---|---|---|---|
| | | **PermaNet® 2.0** | **PermaNet® Dual** | **PermaNet® 3.0** | **PermaNet® Dual** | **Interceptor® G2** | **PermaNet® Dual** |
| | **Total collected** | 1393 | 1255 | 1486 | 1255 | 1292 | 1255 |
| **Mortality (primary endpoint)** | **Total dead** | 241 | 951 | 599 | 951 | 1021 | 951 |
| | **Mortality (%)** | 17.3 | 75.8 | 40.3 | 75.8 | 79.0 | 75.8 |
| | **Odds ratio** | – | 16.000 | – | 5.412 | – | 0.854 |
| | **Std. error (on log odds scale)** | – | 1.612 | – | 0.474 | – | 0.085 |
| | **P-value** | – | <0.001 | – | <0.001 | – | 0.112 |
| | **95% CIs** | – | 13.133–19.492 | – | 4.292–6.160 | – | 0.703–1.038 |
| | **WHO efficacy criteria** | – | Significantly higher (p<0.05) | – | Significantly higher (p<0.05) | – | Lower 95% CI >0.7 |
| | **Conclusion** | – | **Superior** | – | **Superior** | – | **Non-inferior** |
| **Blood-feeding (secondary endpoint)** | **Total blood-fed** | 705 | 433 | 348 | 433 | 339 | 433 |
| | **Blood-feeding (%)** | 50.6 | 34.5 | 23.4 | 34.5 | 26.2 | 34.5 |
| | **Odds ratio** | – | 0.468 | – | 1.519 | – | 1.445 |
| | **Std. error (on log odds scale)** | – | 0.040 | – | 0.141 | – | 0.135 |
| | **P-value** | – | <0.001 | – | <0.001 | – | <0.001 |
| | **95% CIs** | – | 0.395–0.554 | – | 1.267–1.821 | – | 1.203–1.735 |
| | **WHO efficacy criteria** | – | Significantly lower (p<0.05) | – | Significantly lower (p<0.05) | – | Upper 95% CI <1.43 |
| | **Conclusion** | – | **Superior** | – | **Inferior** | – | **Not non-inferior** |

**Table S2:** Supplementary cone bioassay results with the susceptible *Anopheles gambiae sensu stricto* Kisumu strain. *A total of 40–60 mosquitoes were exposed to each of the five net pieces per treatment arm for 3 mins in ten batches of 4–6.*

| Net type | Net status | N | N KD | % KD | 95% CIs | N dead 24 h | % dead 24 h | N dead 48 h | % dead 48 h | N dead 72 h | % dead 72 h | 95% CIs |
|---|---|---|---|---|---|---|---|---|---|---|---|---|
| Control | – | 106 | 0 | 0 | 0-5 | 0 | 0 | 0 | 0 | 0 | 0 | 0-5.0 |
| PermaNet® 2.0 | Unwashed before trial | 55 | 55 | 100 | 95.0-100 | 53 | 96.4 | 55 | 100 | 55 | 100 | 95.0-100 |
| | Washed 20x before trial | 51 | 39 | 76.5 | 64.9-88.1 | 13 | 25.5 | 14 | 27.5 | 19 | 37.3 | 24.0-50.6 |
| | Unwashed after trial | 49 | 41 | 83.7 | 73.4-94.0 | 41 | 83.7 | 41 | 83.7 | 42 | 85.7 | 75.9-95.5 |
| | Washed 20x after trial | 49 | 43 | 87.8 | 78.6-97.0 | 33 | 67.3 | 35 | 71.4 | 36 | 73.5 | 61.1-85.9 |
| PermaNet® 3.0 (sides) | Unwashed before trial | 42 | 39 | 92.9 | 85.1-100 | 38 | 90.5 | 38 | 90.5 | 39 | 92.9 | 85.1-100 |
| | Washed 20x before trial | 42 | 39 | 92.9 | 85.1-100 | 21 | 50.0 | 24 | 57.1 | 26 | 61.9 | 47.2-76.6 |
| | Unwashed after trial | 38 | 36 | 94.7 | 87.6-100 | 35 | 92.1 | 35 | 92.1 | 36 | 94.7 | 87.6-100 |
| | Washed 20x after trial | 45 | 38 | 84.4 | 73.8-95.0 | 40 | 88.9 | 42 | 93.3 | 43 | 95.6 | 89.6-100 |
| PermaNet® 3.0 (roof) | Unwashed before trial | 31 | 27 | 87.1 | 75.3-98.9 | 31 | 100 | 31 | 100 | 31 | 100 | 95.0-100 |
| | Washed 20x before trial | 30 | 30 | 100 | 95.0-100 | 30 | 100 | 30 | 100 | 30 | 100 | 95.0-100 |
| | Unwashed after trial | 28 | 27 | 96.4 | 89.5-100 | 28 | 100 | 28 | 100 | 28 | 100 | 95.0-100 |
| | Washed 20x after trial | 31 | 30 | 96.8 | 90.6-100 | 31 | 100 | 31 | 100 | 31 | 100 | 95.0-100 |
| Interceptor® G2 | Unwashed before trial | 47 | 36 | 76.6 | 64.5-88.7 | 6 | 12.8 | 8 | 17.0 | 8 | 17.0 | 6.3-27.7 |
| | Washed 20x before trial | 50 | 17 | 34 | 20.9-47.1 | 4 | 8.0 | 4 | 8.0 | 7 | 14.0 | 4.4-23.6 |
| | Unwashed after trial | 49 | 32 | 65.3 | 52.0-78.6 | 20 | 40.8 | 21 | 42.9 | 22 | 44.9 | 31.0-58.8 |
| | Washed 20x after trial | 52 | 34 | 65.4 | 52.5-78.3 | 13 | 25.0 | 17 | 32.7 | 17 | 32.7 | 20.0-45.5 |
| PermaNet® Dual | Unwashed before trial | 50 | 45 | 90 | 81.7-98.3 | 22 | 44.0 | 22 | 44.0 | 23 | 46.0 | 32.2-59.8 |
| | Washed 20x before trial | 51 | 27 | 52.9 | 39.2-66.6 | 8 | 15.7 | 10 | 19.6 | 13 | 25.5 | 13.5-37.5 |
| | Unwashed after trial | 52 | 37 | 71.2 | 58.9-83.5 | 24 | 46.2 | 24 | 46.2 | 26 | 50.0 | 36.4-63.6 |
| | Washed 20x after trial | 52 | 32 | 61.5 | 48.3-74.7 | 21 | 40.4 | 26 | 50.0 | 29 | 55.8 | 42.3-69.3 |

**Table S3:** Summary tunnel test results with the pyrethroid-resistant *Anopheles gambiae sensu lato* Covè strain. *A total of 160–240 mosquitoes were exposed overnight to each of two randomly selected net pieces per treatment arm in one replicate tunnel test.*

| Treatment | Wash status | N exposed | Total Fed | N dead imm | N dead 24 h | N dead 48 h | N dead 72 h | N pass | % Imm Mort | % 24 h mort | % 48 h mort | % 72 h mort | 95% CI | % Pass | 95% CI | % Bfd | 95% CI |
|---|---|---|---|---|---|---|---|---|---|---|---|---|---|---|---|---|---|
| Control | N/A | 411 | 299 | 9 | 0 | 3 | 8 | 191 | 2.2 | 2.2 | 2.9 | 4.1 | 2.2-6.1 | 46.5 | 41.7-51.3 | 72.8 | 68.5-77.1 |
| PermaNet® 2.0 | Unwashed before trial | 218 | 44 | 29 | 21 | 38 | 75 | 73 | 13.3 | 22.9 | 30.7 | 47.7 | 41.1-54.3 | 33.5 | 27.2-39.8 | 20.2 | 14.9-25.5 |
| | Washed 20 x before trial | 198 | 69 | 35 | 17 | 30 | 51 | 65 | 17.7 | 26.3 | 32.8 | 43.4 | 36.5-50.3 | 32.8 | 26.3-39.4 | 34.9 | 28.2-41.5 |
| | Unwashed after trial | 211 | 29 | 78 | 25 | 39 | 52 | 70 | 37.0 | 48.8 | 55.5 | 61.6 | 55.1-68.2 | 33.2 | 26.8-39.5 | 13.7 | 9.1-18.4 |
| | Washed 20x after trial | 211 | 55 | 81 | 45 | 53 | 63 | 99 | 38.4 | 59.7 | 63.5 | 68.3 | 62.0-74.5 | 46.9 | 40.2-53.7 | 26.1 | 20.2-32.0 |
| Interceptor® G2 | Unwashed before trial | 202 | 31 | 187 | 13 | 13 | 13 | 42 | 92.6 | 99.0 | 99.0 | 99.0 | 97.6-100 | 20.8 | 15.2-26.4 | 15.4 | 10.4-20.3 |
| | Washed 20 x before trial | 235 | 46 | 212 | 15 | 17 | 18 | 57 | 90.2 | 96.6 | 97.5 | 97.9 | 96.0-99.7 | 24.3 | 18.8-29.7 | 19.6 | 14.5-24.6 |
| | Unwashed after trial | 231 | 23 | 223 | 7 | 7 | 7 | 74 | 96.5 | 99.6 | 99.6 | 99.6 | 98.7-100 | 32.0 | 26.0-38.1 | 10.0 | 6.1-13.8 |
| | Washed 20x after trial | 224 | 40 | 220 | 2 | 2 | 2 | 66 | 98.2 | 99.1 | 99.1 | 99.1 | 97.9-100 | 29.5 | 23.5-35.4 | 17.9 | 12.8-22.9 |
| PermaNet® Dual | Unwashed before trial | 216 | 19 | 188 | 23 | 23 | 23 | 66 | 87.0 | 97.7 | 97.7 | 97.7 | 95.7-99.7 | 30.6 | 24.4-36.7 | 8.8 | 5.0-12.6 |
| | Washed 20 x before trial | 224 | 15 | 215 | 8 | 8 | 8 | 53 | 96.0 | 99.6 | 99.6 | 99.6 | 98.7-100 | 23.7 | 18.1-29.2 | 6.7 | 3.4-10.0 |
| | Unwashed after trial | 229 | 11 | 227 | 2 | 2 | 2 | 47 | 99.1 | 100 | 100 | 100 | – | 20.5 | 15.3-25.8 | 4.8 | 2.0-7.56 |
| | Washed 20x after trial | 238 | 17 | 229 | 6 | 6 | 6 | 69 | 96.2 | 98.7 | 98.7 | 98.7 | 97.3-100 | 29.0 | 23.2-34.8 | 7.1 | 3.9-10.4 |
